## Supplementary Material for "PrimPol-dependent single-stranded gap formation mediates homologous recombination at bulky DNA adducts"

### Supplementary Figures

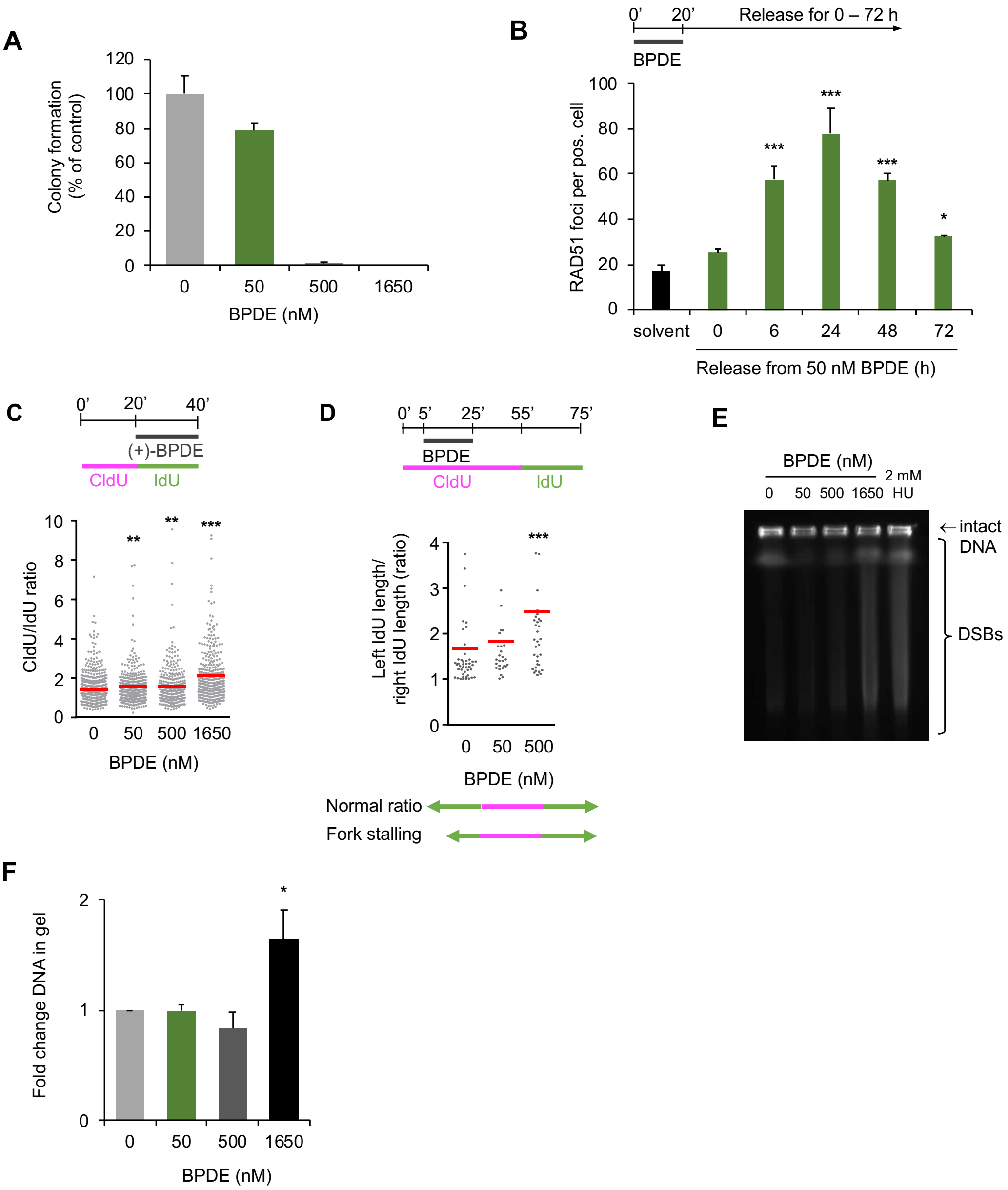


**Fig S1. Replication stress induced by different BPDE concentrations.** (A) Colony survival of U2OS cells after release from 20 min BPDE. n=1 (B) Numbers of RAD51 foci per RAD51 foci positive cell after release from 50 nM BPDE. n=3 (C) CldU/IdU ratios after DNA fibre assay with BPDE present during the IdU pulse. Increased CldU/IdU ratios indicate BPDE-induced fork slowing. (D) Fork asymmetry of bidirectional replication forks after DNA fibre assay with BPDE. Increased left IdU/right IdU ratios indicate fork stalling. (E) Pulse-field gel electrophoresis (PFGE) to visualise DSBs in after 3 hours release from BPDE. Treatment with 2 mM HU for 48 hours was included as a positive control. (F) Quantification of PFGE. DSBs were normalized to the total amount of DNA, then normalised to control. n= 5. The means and SEM (bars) of independent experiments are shown. Asterisks indicate p-values compared to control unless indicated otherwise (one-way ANOVA for B, F, Mann-Whitney for C, D, * $p < 0.05$, ** $p < 0.01$, *** $p < 0.001$)

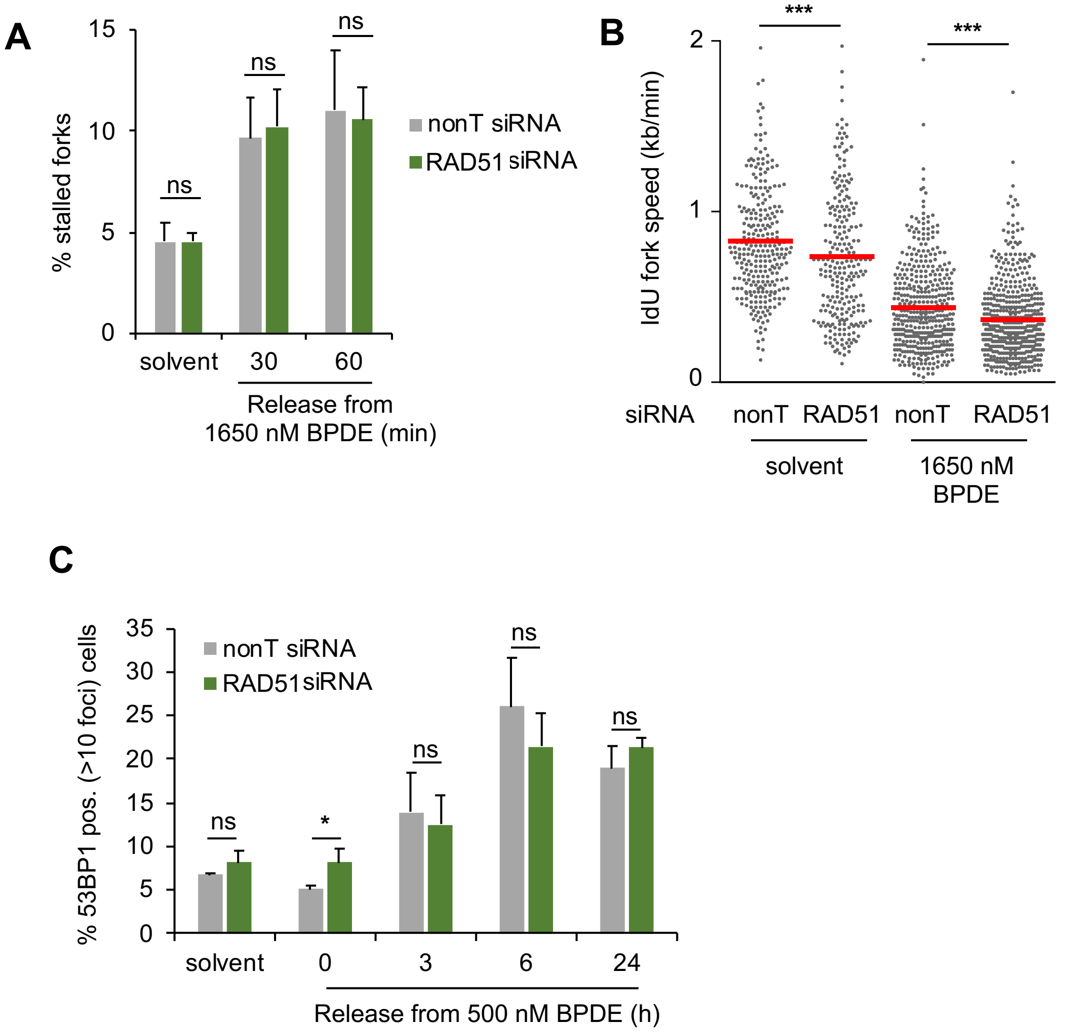


**Fig S2. HR induced by different BPDE concentrations.**
(A) Quantification of stalled forks after release from 1650 nM BPDE for 30 or 60 min in cells treated with nonT or RAD51 siRNA. n=3 (B) Replication fork speeds in cells 30 min released from treatment with 1650 nM BPDE in presence or absence of RAD51 siRNA. (C) Percentages of control- or RAD51-depleted U2OS cells with > 10 53BP1 foci after release from 500 nM BPDE. n=3. The means and SEM (bars) of independent experiments are shown. Asterisks indicate p-values (student's t-test for A, C or Mann-Whitney for B, * $p < 0.05$, *** $p < 0.001$).

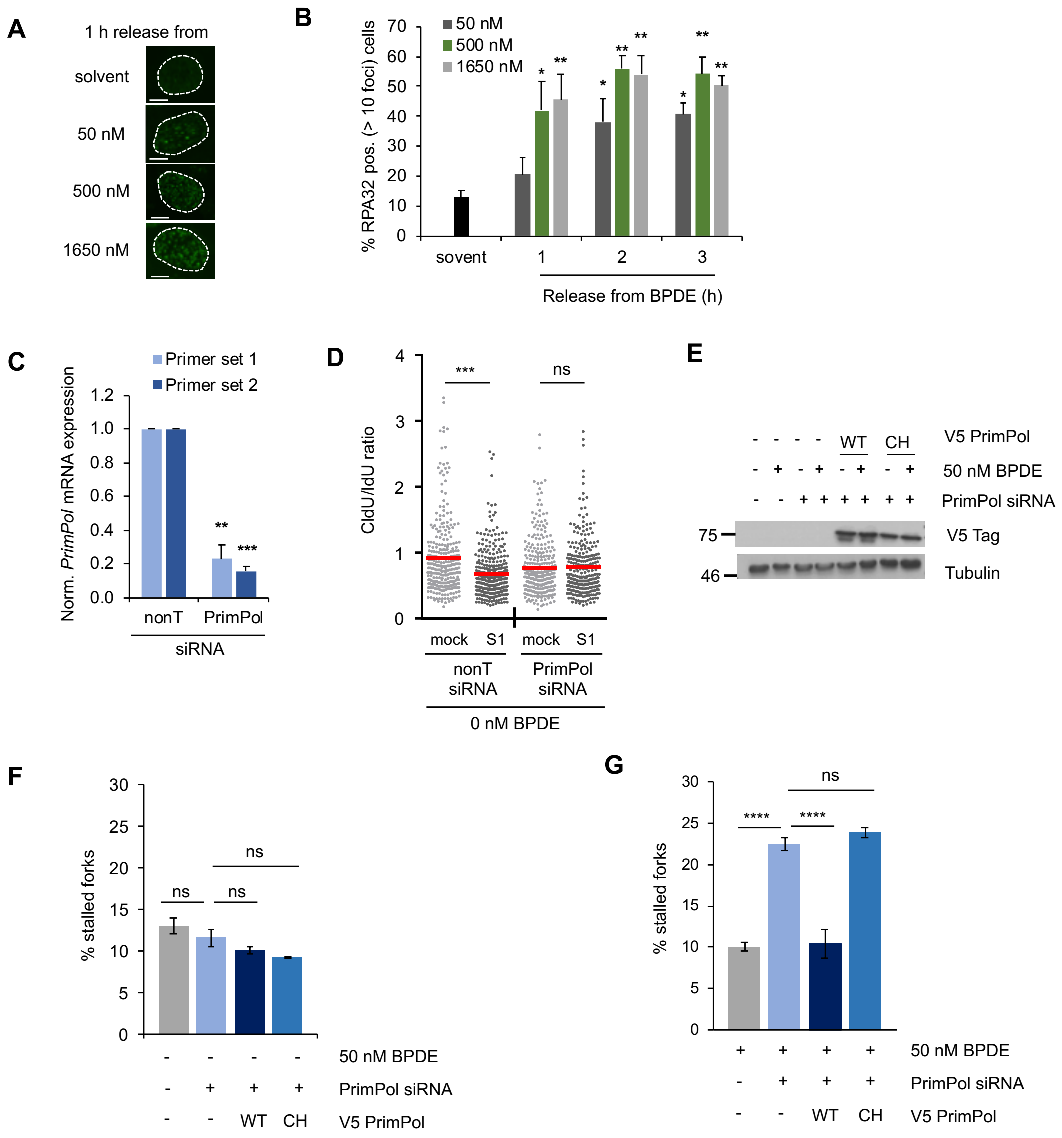


**Fig S3. BPDE induces single-stranded DNA gaps via PrimPol-dependent re-priming.**

(A) Representative images of RPA foci after 1 h release from 20 min BPDE as indicated. (B) Percentages of U2OS cells with > 10 RPA foci after release from BPDE for the times indicated. n=3 (C) *PRIMPOL* mRNA quantification by qRT-PCR after 48 h siRNA transfection. *PRIMPOL* mRNA levels were normalised to *RPLP0* and control. n=3 (D) CldU/IdU ratios after S1-modified DNA fibre assay in cells treated with solvent as in Fig. 2G after 48 h of control or PrimPol siRNA. n=3 (E) Protein levels of V5-tagged WT or CH PrimPol and Tubulin (loading control) after release from 50 nM BPDE in presence of nonT (-) or PrimPol siRNA (F) Quantification of stalled forks after release from solvent in cells treated with nonT or PrimPol siRNA and expression constructs encoding GFP (-) or siRNA-resistant WT or CH PrimPol. n=3. (G) Quantification of stalled forks after release from 50 nM BPDE in cells treated as in F. n=3. The means and SEM (bars) of independent experiments are shown. Asterisks indicate p-values compared to control (one-way ANOVA for B, C, F, G and Mann-Whitney for D, * $p < 0.05$, ** $p < 0.01$, *** $p < 0.001$).

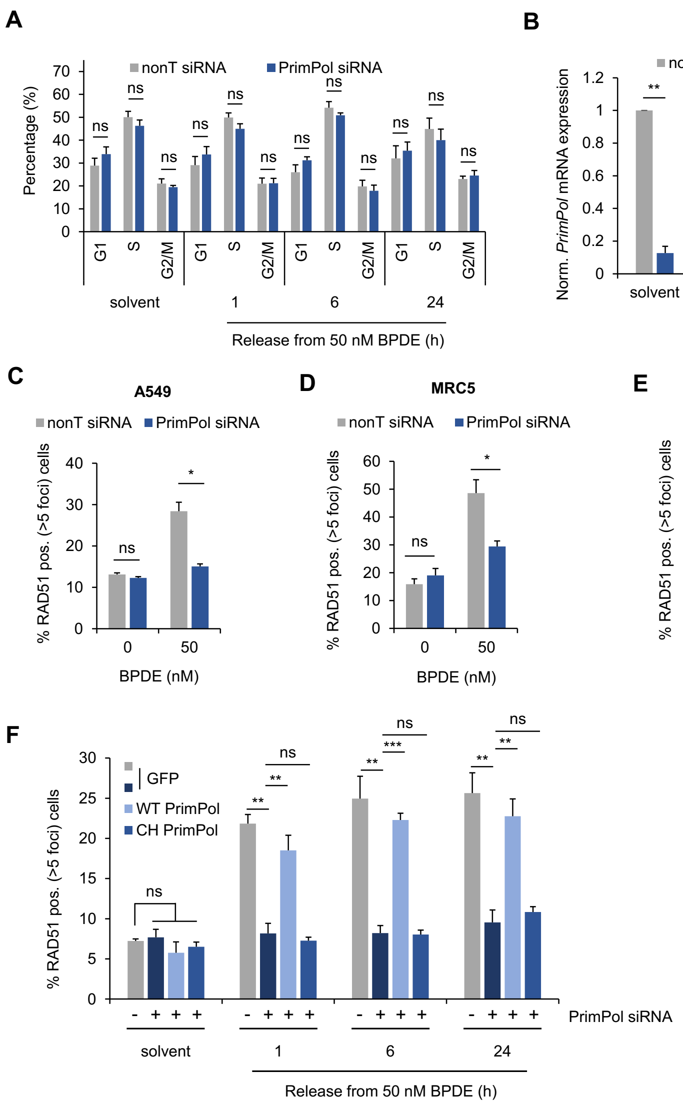


**Fig. S4. Impact of PrimPol depletion on cell cycle distribution, and on RAD51 loading in response to BPDE in different cell lines.**
(A) Quantification of cell cycle distributions after release from solvent (1 h) or 50 nM BPDE in presence of nonT or PrimPol siRNA, as in Fig. 3C. Cells were labelled with BrdU for 30 min to label replicating cells prior to flow cytometry. n=3. (B) *PRIMPOL* mRNA quantification by qRT-PCR after 48 h siRNA transfection and release from 20 min 50 nM BPDE as indicated. *PRIMPOL* mRNA levels were normalised to *RPLP0*, non-treated control, and the respective solvent-treated controls for each time point. n=3. (C, D) Percentages of A549 (C) or MRC5 (D) cells with >5 RAD51 foci after 1 h release from 50 nM BPDE, in presence of nonT or PrimPol siRNA. n=3. (E) Percentages of MRC5 WT or PrimPol $^{-/-}$ cells with >5 RAD51 foci after 1 h release from 50 nM BPDE. n=3. (F) Percentages of U2OS cells with >5 RAD51 foci after release from 50 nM BPDE in presence of nonT (-) or PrimPol siRNA and expression constructs encoding GFP (-) or siRNA-resistant WT or CH PrimPol. n=3. The means and SEM (bars) of independent experiments are shown. Asterisks indicate p-values as indicated (student's t-test for A, B, C, D, E or one-way ANOVA for F * $p < 0.05$, ** $p < 0.01$, *** $p < 0.001$).

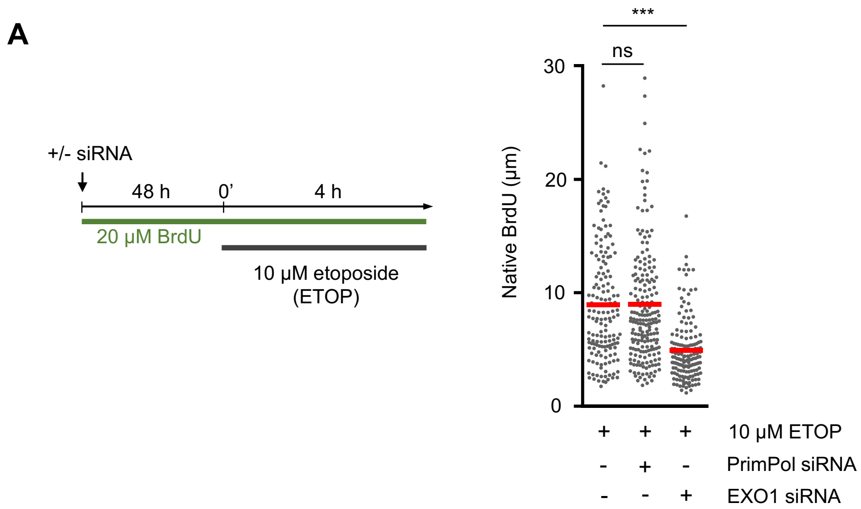


**Fig. S5. ssDNA track lengths at etoposide-mediated DSBs are independent of PrimPol.**
(A) SMART analysis of single-stranded DNA after treatment with etoposide as indicated, in presence of nonT (-), PrimPol, or EXO1 siRNA for 48 h. The bars show means of one independent experiment. Asterisks indicate p-values (Mann-Whitney, *** $p < 0.001$).

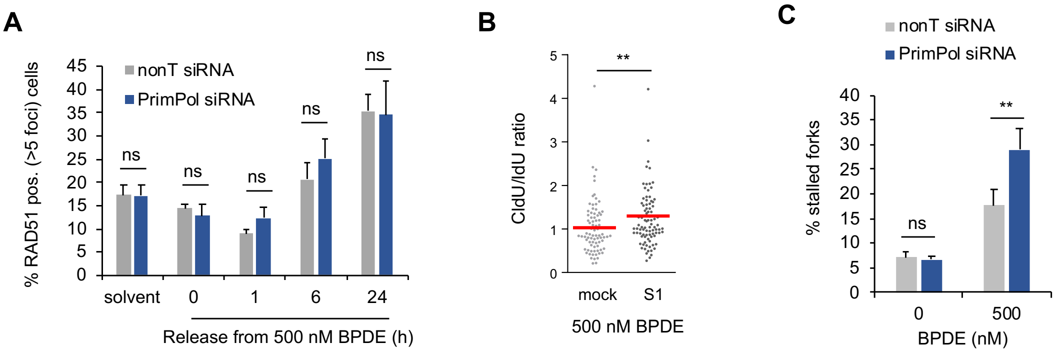


**Fig. S6. PrimPol mediates RAD51 loading in response to replication transcription conflicts induced by JQ1, but not at DSBs induced by high BPDE concentrations.**
(A) Percentages of cells with >5 foci after release from 500 nM BPDE for the times indicated, in presence of nonT or PrimPol siRNA. n=3 (B) CldU/IdU ratios after S1-modified DNA fibre assay in cells treated with 500 nM BPDE as in Fig. 2B. (C) Quantification of stalled forks after release from 500 nM BPDE in presence of non-targeting (nonT) or PrimPol siRNA. Fibre labelling was according to Fig. 2F. n=3. The means and SEM (bars) of independent experiments are shown. Asterisks indicate p-values (student's t-test for A, C and Mann-Whitney for B, ** $p < 0.01$, *** $p < 0.001$).

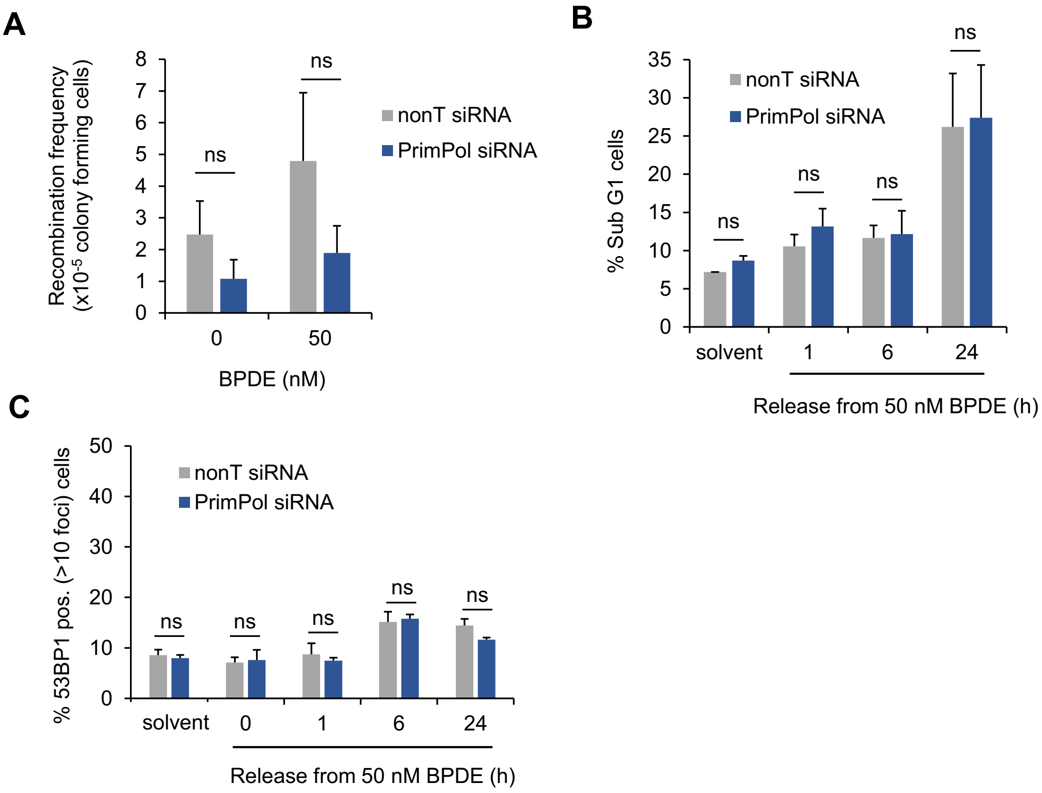


**Fig. S7. Impact of PrimPol on recombination, cell death and DSB formation induced by bulky DNA adducts.** (A) Absolute recombination frequencies in SW480SN.3 cells induced by 50 nM BPDE in presence of nonT or PrimPol siRNA. n=3-4 (B) FACS analysis of percentages of sub-G1 cells after release from solvent or 50 nM BPDE in presence of nonT or PrimPol siRNA. Release from solvent was for 1 hour. n=3-4 (C) Percentages of control- or PrimPol-depleted U2OS cells with > 10 53BP1 foci after release from 50 nM BPDE. n=3**.** The means and SEM (bars) of independent experiments are shown. Non-significant differences between samples were determined by student's t-test.

**Supplementary Table S1. Sequences of qRT-PCR primers**

| **Name** | **Sequence (5' → 3')** |
|---|---|
| **PRIMPOL1 For** | CCGAGGTATCCCAGAGGTGA |
| **PRIMPOL1 Rev** | AATGCCCCACGTTGCTTTTC |
| **PRIMPOL2 For** | AAAAGCAACGTGGGGCATTC |
| **PRIMPOL2 Rev** | GGTGGTTCTTCTGGCTTGGA |
| **RPLP0 For (control)** | CAGATTGGCTACCCAACTGTT |
| **RPLP0 Rev (control)** | GGAAGGTGTAATCCGTCTCCAC |
